# A new paradigm for biological sequence retrieval inspired by natural language processing and database research

**DOI:** 10.1101/2023.11.07.565984

**Authors:** Axel-Jan Rousseau, Sébastien Lemal, Yegor Korovin, Georgios Triantopoulos, Ingrid Brands, Maxim Biemans, Dirk Van Hyfte, Dirk Valkenborg

## Abstract

Nearly-exponential growth and heterogeneity of biological sequence data make the task of biological sequence retrieval from databases more important and challenging than ever. In this manuscript, we present a novel search algorithm involving an indexing scheme based on patterns discovered by natural language processing, i.e., short strings of nucleotides or amino acids, akin to standard k-mers, but mined from cumulative cross-species omic data repositories. More specifically, we benchmark the quality of the sequence retrieval process by comparing to BLASTP, a heuristic algorithm for the alignment of genomics or protein sequence data. The main argumentation is that to retrieve biological similar sequences it is not needed to mimic the alignment procedures as it is performed by BLAST. Our results suggests that the HYFT-indexing and searching is a good alternative and a static, alignment-free method to retrieve homologous sequence down to 50% sequence identity.

## Introduction

Life science researchers and molecular biologists face an ever growing amount of biological sequence information [1–3] that is embedded in a plethora of public repositories (EBI [4], NCBI [5], ELIXIR [6], UNIPROT [7], REFSEQ [8], Genbank [9], MASSBank [10], *etc*.). These databases also serve different clinical purposes (cancer, population variation, immunodeficiency, allergens) across the different molecular layers (DNA, RNA, AA) and are often specialized towards certain target areas (immunoglobulins, t-cell receptor repertoire, major histocompatibility complexes), or only contain partial sequence information on relevant subsets, like antigen, domains, motifs, or pattern databases [11]. Considering the scale and heterogeneity of these information repositories, they pose a problem for database retrieval systems that today rely on the Basic Local Alignment Search Tool (BLAST) or BLAST-like searches or their derivatives [12].

BLAST is an ingenuous system, but we argue that with the exponential growth of biological sequence information and the high-diversity of current public databases, it is of crucial interest to investigate other principles to devise high-performing retrieval systems that are better suited to cope with drastic data growth, and present a more holistic selection of results across the different types of molecular layers. To tackle the problem of homology search in the ever increasing amount of data, alignment-based methods using suffix arrays such as GHOSTX [13] and RAPSearch [14] and indexing such as DIAMOND [15, 16] have been proposed.

In this manuscript, we want to explore alternative paths for biological sequence retrieval that are partially inspired by the information present in pattern-databases like PRINTS [17] and PROSITE-PATTERN [18]. The general idea is that these well-conserved patterns with a length between five and eight, could be used as anchor points and prior information for indexing purposes. The requirement for this idea to work is that many of these anchor points exist and that they are densely and uniformly distributed over the biological sequence database. Unfortunately, the number of patterns in the aforementioned databases are limited to only a few thousand patterns and they are too sparsely distributed over the biological sequence databases.

However, the leverage comes from the field of computer science and database research, where we employed algorithms from natural language processing (NLP) to generate novel and meaningful patterns from publicly available databases, in analogy with to the bioinformatics tool PRATT [19]. In the remainder of this manuscript, these newly discovered patterns are termed HYFT patterns [20]. A description of these patterns is given in Sec. HYFT patterns.

At this point, it is worthwhile to reiterate that the objective of this research is not to develop a new sequence alignment method, but rather a static sequence retrieval method. With a static algorithm we indicate a procedure that does not rely on dynamic programming, i.e., alignment or any other dynamic programming action like, e.g., gap extension. Hence, a static search solely relies on a database look-up, and any innovations in database technology will also benefit HYFT-based search, as these HYFT patterns are indexed prior to any queries.

The current trend is to use alignment-free sequence comparison methods, many of wich rely on statistics derived from k-mers [21] and [22]. These methods do not perform an alignment between the query and database-sequences, but instead calculate a distance metric to measure similarity between sequences. Our newly proposed method based on HYFTs, however, does not calculate distances, nor does it make use of k-mers.

HYFT patterns, which are enriched with various metadata and which we describe in more details in Sec. HYFT patterns, can be used for database indexing irrespective of the nature of the sequences (nucleotides or amino acids). This step will result in fast data retrieval and increased scalability as the indexing procedure has to be performed only at the initialization of the database. In order to investigate whether a particular query sequence of interest is present in that database, it will be deconstructed into a corresponding sequence of HYFT patterns. Next, all the sequences that link to the indexed HYFT patterns in the query are retrieved and further filtered for confidence.

In order to evaluate the HYFT-search algorithm, a study to benchmark the quality of sequence retrieval was set-up similar in spirit to previous studies [23–26], where we searched the Protein Data Bank [27, 28] (PDB) for structural homologous sequences facilitated by PDB that also hosts the structural models associated to these proteins sequences. As a ground truth dataset, we considered the CATH [29] database which assigns to PDB entries a structural classification label (for example model 101M, chain A [30, 31] is annotated with 1.10.490.10) and we calculated truncated Receiver Operator Characteristic (ROC) and Precision-Recall curves to compare the HYFT-based search with a BLASTP search. The latest version of blastp is chosen because this is still the most commonly used algorithm.

The scope of this investigation is to evaluate the sensitivity and specificity achieved with BLASTP versus the HYFT-based alignment-free sequence retrieval algorithm. Other characteristics that might be of interest in comparing algorithm performance such as speed of retrieval or scalability were not considered in this manuscript.

This manuscript is organized as follows: first, we detail the methodology, then present the results. Following this, we discuss the specificities of each approach and how they impact the results. Finally, we discuss how the HYFT-search can be improved, why HYFTs can be used in tasks requiring highly related homologs, and how HYFTs are of interest for new scientific discoveries in the context of these applications.

## Materials and methods

In this section we give an overview of the HYFT technology. In the first section we explain the HYFT patterns, next we discuss how the HYFTs are used to index large biological databases and we are specific about the underlying technologies used. We also explain how the query sequence is parsed and related to the HYFT database, as not all possible HYFT patterns are used for searching. The search algorithm is also detailed. Some post-processing of the results is also described in this section. In the two concluding sections, we summarize the BLAST algorithm to contextualize the difference with the HYFT approach, we discuss the HYFT statistics and figures of merits used to conduct this bench-marking. It should be cautiously noticed that only for bench-marking purposes and the construction of a fine-grained ROC curve, the sequences retrieved by the HYFT patterns were aligned using the Smith-Waterman score. The alignment step is not included in the HYFT algorithm but only performed to be able to compare the HYFT retrieval with the BLAST e-values.

### HYFT patterns

In the implementation used in this paper we used HYFT patterns ranging from a length of 5 amino acids up to a length of 7 amino acids. In typical *k*-mer based approaches, the size *k* of the *k*-mer is fixed, and all *k*-mers found in sequences can be used for the look-up table. In some k-mer based implementations filtering steps such as removing low-complexity *k*-mers, or applying a substitution matrix score threshold are used to reduce the number of look-ups.

The core idea behind the HYFT methodology is to apply NLP methods to sequence data. NLP is a sub-field of linguistics, computer science, and artificial intelligence concerned with building tools and algorithms to help computers to process textual data written in a natural language (i.e. used by human beings for communication). A specific application of NLP is the identification of tokens, semantic units of text carrying conceptually relevant information.

In analogy to NLP, HYFTs [20] are to biological sequences what tokens are to textual data. A single HYFT is a data object encompassing contextual information (i.e. beyond the associated pattern itself). HYFTs are identified by mining aggregated biological sequence databases spanning the full known biome for patterns, which are short enough so that they are less prone to indels or substitutions, specific enough to characterise the parent sequence with respect to factual knowledge; moreover, there must be a sufficient amount of these patterns so that they can be retrieved from any sequence (hence their distribution within the data must be dense enough in order to be used for indexing). They are unique signature sequences, akin to biological fingerprints, in AA, but also DNA and RNA. With HYFTs, all biological data, irrespective of species or function, can be tokenized (i.e. divided into sub-units) to a common omics language. This contrasts with standard *k*-mer (substrings of sequence of a fixed length *k*) approaches, which typically do not encompass contextual information. In the following, we refer to “specificity” as a feature of the HYFTs with respect to context in their direct neighbourhood, which allows us to rank them according to this criterion. HYFTs strike a good balance between the high specificity of long patterns and the lower specificity of short patterns.

HYFTs are fundamentally different from usual functional patterns or motifs afore-mentioned in the introduction, as they are not defined in terms of biological priors. HYFTs are mined from any string-like data structures.

HYFTs have the following features: i) the pattern length is not fixed, but is usually small; ii) an integer is associated to the pattern which encodes the different occurrences of amino acids in the vicinity of the pattern and iii) a rank (integer) is associated to each HYFT, designed so that HYFTs with small combinatorial numbers have a lower rank. This defines the aforementioned “specificity” feature.

### Parsing and database indexing

Now that we have a database with HYFT patterns, we need to parse these HYFT patterns and index these patterns on the entire biosphere of biological databases to obtain a look-up table. The parsing step concerns the retrieval of a limited number of HYFTs from a database sequence, as not all HYFTs are necessarily relevant. This is similar in spirit as the usage of the DUST filter in BLAST where low informative regions are removed from seeding. The indexing part is about building a look-up table based on the HYFTs retrieved from the parsing step.

#### Parsing

A limited number of HYFTs retrieved from a sequence are used for indexing. In the current approach, which we will refer to as specific or non-overlapping, the HYFT parser prioritizes those with the lower rank which do not overlap. First, all HYFTs that occur are enumerated. Then, the parser select the most specific one (lowest rank) which do not overlap (if two overlap, the one with the lower rank is selected). This process is repeated for the region between the HYFT, iteratively, until no more HYFTs can be found: this limits the number of HYFTs extracted from a sequence, and ensure that most of the sequence is covered by HYFTs. The HYFT length (or word size) is not a parameter, in contrast to BLAST. This parsing step is then used for indexing and for querying/matching.

#### Indexing

The parsing step ultimately limits the number of HYFTs retrieved from a sequence. The index is constructed as a standard look-up table: all sequences are parsed, and the HYFTs retrieved are stored paired with their corresponding sequence identifier. Because not all HYFTs from a sequence are used, the size of the index is ultimately limited. The average number of HYFTs per sequence length is 0.12. In contrast, the theoretical upper bound on the distinct *k*-mers retrieved from a sequence of length *l* is *l* − *k* + 1 (assuming no repetition of the *k*-mer within the sequence) and the ratio *l* − 1 + *k/l* is close to 1. Hence, there is approximately a factor of 8 reduction in terms of extracted patterns between HYFTs and *k*-mers of similar size.

The index and sequence metadata used in this benchmark are stored in the Inter-Systems IRIS data platform, deployed on an Amazon Elastic Compute Cloud (EC2) instance with the r5.xlarge instance type configuration (with 4 Intel Xeon Platinum 8000 CPUs and 32 GiB of memory [32]). The communication between the database and a user is ensured by a RESTful API exposed by the IRIS database.

### Query parsing and look-up

With the biological databases fully indexed a database query becomes convenient, scalable and fast. For each query sequence, the putative HYFT patterns are filtered from the sequence - a step that is explained in the next subsection. Once the relevant HYFT patterns are known via a look-up table the database sequences are retrieved - see subsection Look-up;

#### Query parsing

When a query sequence is submitted, it is parsed with the same logic as the sequences in the database used in the indexing approach, resulting in an array of non-overlapping, specific HYFTs.

#### Look-up

For sequence retrieval, matches are retrieved by looking up the sequence identifiers associated with the identified HYFTs in the query sequence. A match is conserved if a sequence in the database contains at least 2 HYFTs (or 1 for short queries with from which a single HYFT is extracted) in common with the query sequence. All such sequence are retrieved and considered as related sequences to the query. In the HYFT approach, this list is prioritized by the number of HYFTs per sequence. In this bench-marking approach a fine-grained score is computed by aligning the query to the returned sequences by Smith-Waterman as explained in Sec. Set-up of the comparison between HYFT pattern search and BLAST.

### BLAST search

The BLAST algorithm also makes use of a look-up table, albeit an index of the query sequence, and it uses *k*-mers for indexing. The word size can be set to different values (from 3 to 6 amino acids). The database sequences are then linearly scanned for *k*-mers in the query. In addition to this look-up, BLAST then extends a hit to find a longer alignment pair between the query and sequence. This extension continues until the alignment score drops below a certain threshold [12]. Then, high scoring pairs are further aligned using ungapped alignment. The alignment process is the dynamic component within the algorithm that goes beyond a pure static index-based retrieval.

### HYFT statistics

The mean coverage, defined as the sum of the length of non-overlapping HYFTs divided by sequence length, is 83% with respect to full sequence using the PDB dataset purged of redundant sequences (at 100% identity level), as shown in Fig. 1.a, b and c. There are outliers to the main distribution, mainly due to sequence models with missing amino acids. These regions are annotated with “X”s, and no HYFT pattern can be found there. The regions in between HYFT patterns (defined as interdistance) have a size ranging from 0 to 6 amino-acids, the distribution is plotted in Fig. 1.d. The distribution of the number of HYFTs per sequence in the indexed database is shown in Fig. 1.e: the average number of HYFTs is roughly 33, after the third peak at this value, the frequency rapidly decreases. This distribution follows that of the sequence lengths in PDB, consistently with the coverage distribution (Fig. 1.c). Finally, in Figure 1.f, the number of HYFTs obtained with the parsing scheme described in Sec. is plotted versus the total number of overlapping HYFTs possibly retrieved, for each sequence: it follows a linear trend, with a slope of ∼ 13. A look-up table built with the specific, non-overlapping scheme is thus on average 13 times smaller than a table built using all possible HYFT patterns.

**Fig 1.**
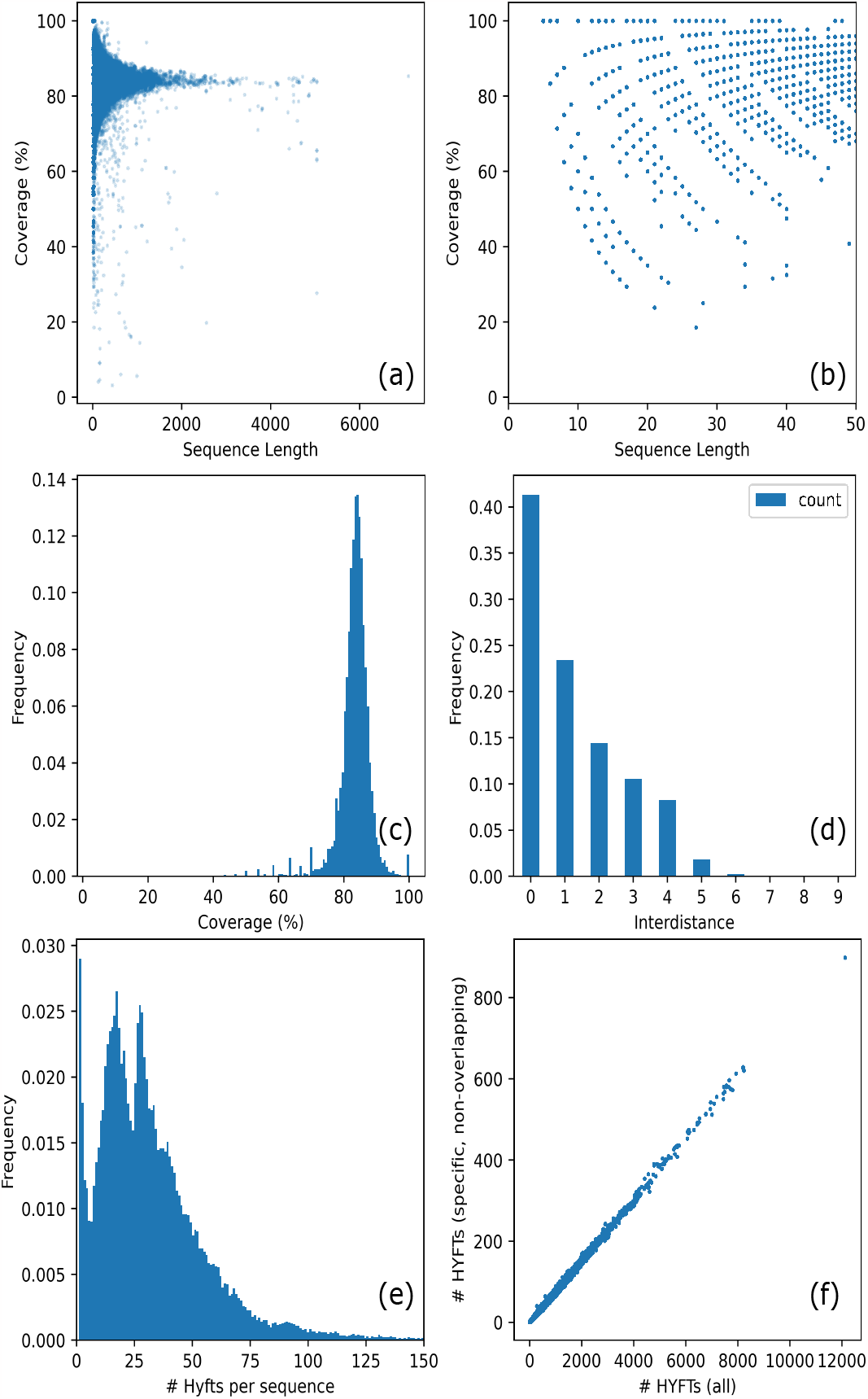
Statistics of HYFTs retrieved from PDB: (a) coverage of HYFT patterns with respect to full sequence; (b) a zoomed-in coverage of HYFT patterns for sequence with length *<* 50; (c) distribution of HYFT coverage; (d) distribution of distance in-between HYFT; (e) distribution of the number of HYFT per sequence and (f) number of specific, non-overlapping HYFTs versus all overlapping HYFTs retrieved in sequences.

### Set-up of the comparison between HYFT pattern search and BLAST

To compare the HYFT pattern search with BLAST, we benchmarked its performance in a similar fashion as in Refs. [33, 34]. The CATH database was used as a ground truth for protein similarity. This database contains protein domains from PDB polypeptide chains, and classifies them into a structural hierarchy based on their sequence and structure. When there is sufficient evidence that domains are related by evolution, they are classified into the same Homologous Superfamily group.

We randomly sampled one domain from each of the 100 largest superfamilies, and queried these against PDB. The PDB chain sequences can contain multiple domains, and can therefore be associated with multiple CATH classifications. For each query, we counted a result from the search as relevant (true positive, or TP) when one of the chain’s superfamily matched the query domain’s class, and as irrelevant (false positive, or FP) when the chain only contained domains of different CATH superfamilies. The unprocessed fasta file for the PDB data can contain multiple duplicate sequences with different identifiers. Furthermore, not all PDB sequences are covered by the CATH database. We therefore only considered PDB chains with a CATH classification, and collected class info based on unique sequences of chains. An overview of the benchmarking workflow is shown in Figure 2)

**Fig 2.**
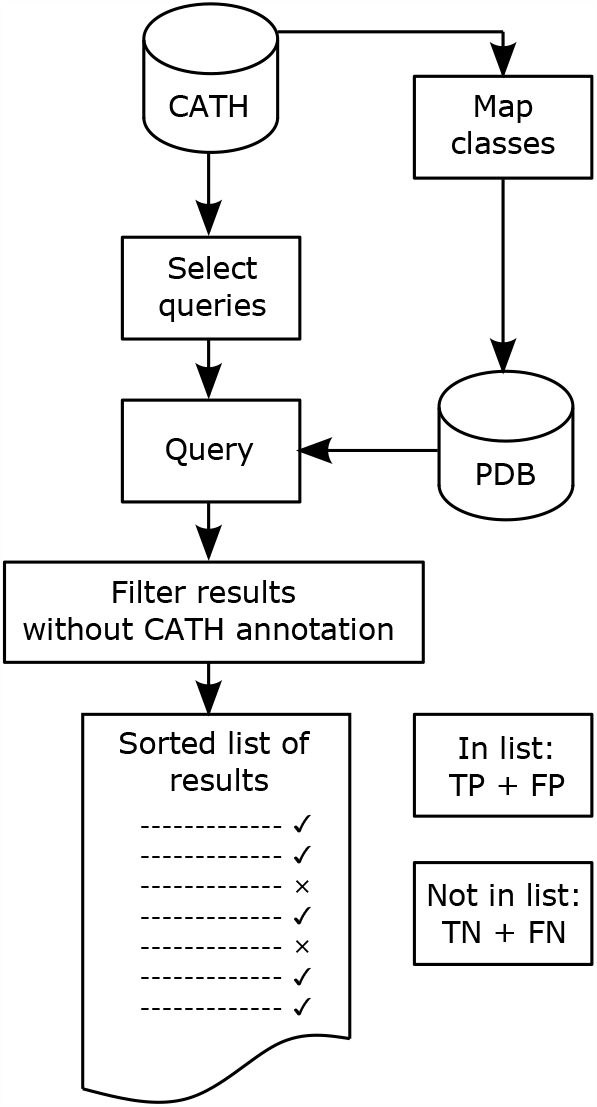
Benchmarking workflow of the present study. Since not all PDB sequences are annotated in CATH, and since not all CATH sequences are in PDB, the classes of CATH domains are mapped back to the PDB sequences the domain belonged to. Then, 100 domain queries are selected from the largest CATH superfamily classes, and are queried against the PDB dataset. Some sequences can have multiple domains belonging to different classes, and others have no domain classified in CATH. These are filtered out. The result is a list of sequences that the search algorithm deems positive, or relevant to the query.

The results are compared with a BLASTP (the BLAST flavor for protein search) search of the PDB database created from the fasta files available from the RCSB downloads page [35]. Default parameters are used for BLASTP [36], except for the `−max_target_seq` (maximum number of aligned sequences to keep) which was set to the size of the database, as otherwise the search would stop early due to the many duplicates found.

### Figures of Merit

The protein chains returned by the search algorithm are considered to be positive matches, those not returned are considered to be negative. These two sets were further divided into true positives when one of the chain’s superfamily matched the query domain’s class, and false positives when it did not match. Using the true positives, false positives, and the total amount of positives and negatives, we defined metrics to assess the quality of the search. The Precision or positive predictive value is the fraction of true positives among the retrieved sequences:

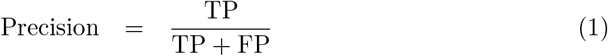

The Recall, also known as the True Positive Rate (TPR) is the fraction of positives retrieved

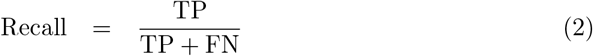

The False Positive Rate (FPR) is the fraction of negative sequences incorrectly identified as positive:

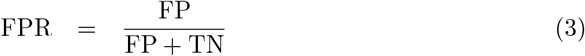

Precision, Recall and FPR are illustrated in Figure 3.

**Fig 3.**
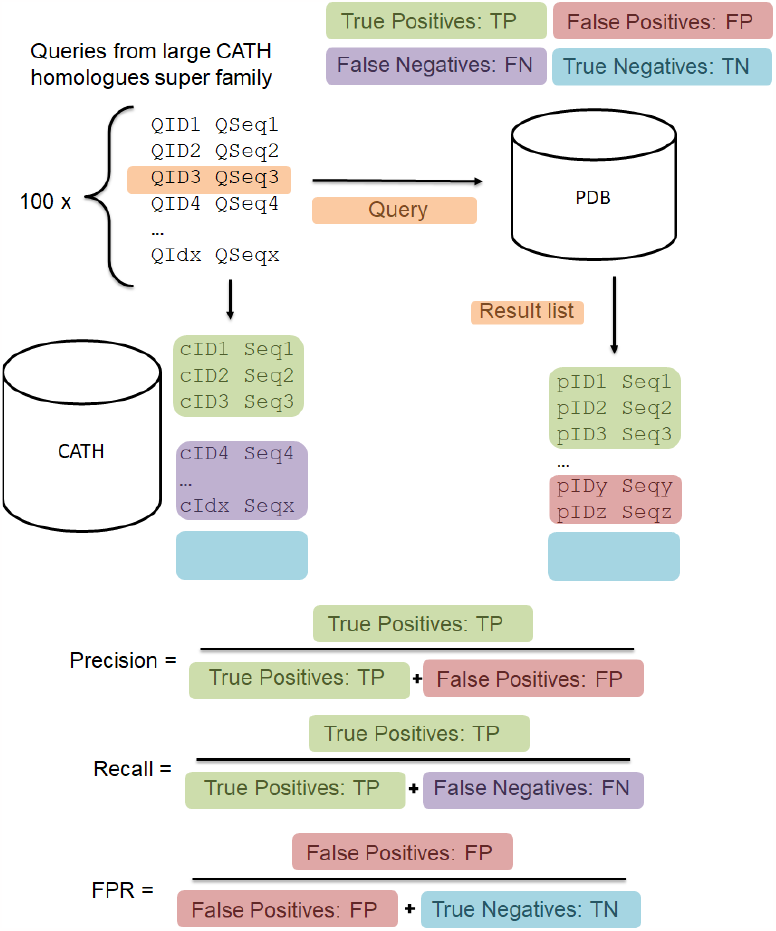
Diagram representation of true positives, false positives, false negatives, true negatives, Precision, Recall (or True Positive Rate) and False Positive Rate.

These metrics were combined to calculate different Figures of Merit characterising the performance of the search task. The Receiver Operating Characteristic (ROC) curve is a commonly used tool to investigate the performance of binary classifiers [37, 38]. For the BLASTP search, the list of search results is ordered by e-value; this is number of expected hits of similar quality score that could be found randomly in a database of a given size: the lower the e-value, the more likely the hit is a relevant match. For the HYFT-search, a local alignment score is used to sort the results. This alignment is not part of the HYFT retrieval algorithm but was used as a way to rank results from different queries in a consistent manner. The Recall is plotted against the FPR by going through the result list at different thresholds. A perfect classifier would rank every TP above the FP, placing the plot in the upper left. A random ordering of the results plots the identity. The area under the ROC curve is the ROC score or AUC, and indicates the probability that a randomly chosen positive is ranked above a random negative.

A complete ROC curve would require that all the sequences in the search database are ranked, but search algorithms commonly return only the fraction of the data deemed relevant. Therefore, our plots show the curve up until the last result returned by the search method is reached. Furthermore, researchers are unlikely to go through several pages of irrelevant results. Therefore, a modification of the ROC is often used: The ROC_*n*_ is the area under the ROC curve plotted until *n* negatives are retrieved, and is a commonly used metric for retrieval in bioinformatics [39]. A value of *n* = 50 is typically used.

In information retrieval the Precision-Recall curve, in which precision is plotted against recall, is more commonly used because it is more suited for tasks with a large class imbalance [40–42]. The precision indicates the fraction of sequences retrieved that are related to the query, which is more relevant in the case where the positive class is given more importance. The area under the precision-recall curve is the average precision (AP):

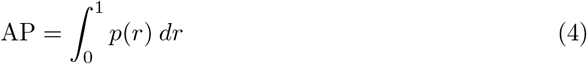

where *p*(*r*) is the precision with respect to recall. The average precision of a set of queries is calculated by taking the mean AP.

## Results

We are now presenting the results of our benchmarking study. In addition to the algorithm described in Section, we also discuss an prototypical improvement of the algorithm involving HYFT extension.

### Benchmarking

Hundred queries were run and results were classified as TP or FP with respect to query class. When searching for homolog superfamilies in PDB, the 100 query domains resulted in a dataset containing 42, 813 positive pairs, and 6, 961, 087 negative pairs. Results are shown in the ROC curve and precision-recall curves in Figs. 5 and 4. In both figures, results for all queries are pooled together and treated as a single sorted result. Both plots show a similar picture. The precision, or fraction of relevant sequences in the returned results, and the FPR of both methods, remain similar until a recall of just over 0.10, meaning 10% of all related sequences were returned. After this point, the HYFT-search algorithm returns more false positives causing the FPR to increase rapidly and the precision to drop, while BLASTP continues to find more true positives up to a recall of 29.4%.

**Fig 4.**
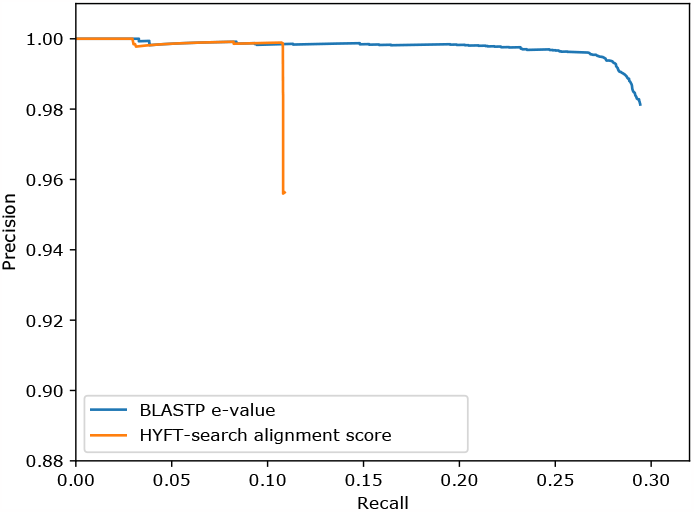
Precision-recall curve for BLASTP and HYFT-search, using the e-value (BLASTP) and the alignment score (HYFT-search) for sorting the results.

**Fig 5.**
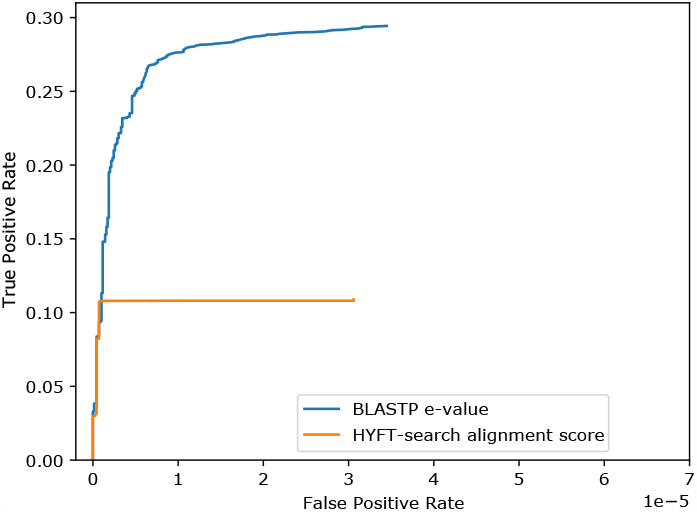
ROC curve for BLASTP and HYFT-search, using the e-value (BLASTP) and the alignment score (HYFT-search) for sorting the results.

The higher Recall for BLAST indicates that more of the sequences deemed relevant are found. To examine the properties of the true positives found by HYFT-search, we compared the overlap between the results returned with these of BLASTP and grouped them by their percent identity to plot the histogram shown in Fig. 6. The histogram shows that a large proportion of the extra TP found by BLASTP comes from pairs with low percent identity. For the pairs with a percent identity above 65%, the HYFT-search is able to retrieve almost the same sequences as BLASTP.

**Fig 6.**
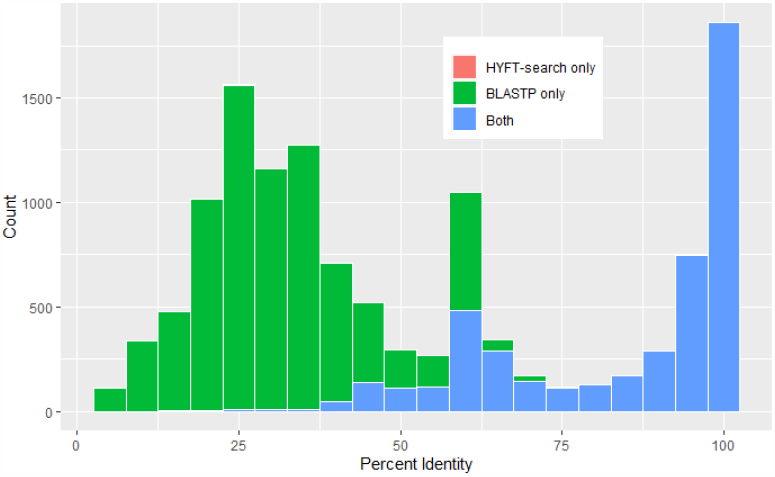
Distribution of True Positives with respect to identity score.

A significant proportion of expected hits are found below 50% identity score. BLASTP performs better in retrieving sequences with an identity score lower than 50%. In order to increase the recall of HYFT-search, we have experimented with a word substitution algorithm similar to that of BLAST, where new HYFTs, not originally found in the query sequences are introduced. These new HYFTs are pre-computed HYFTs that are syntactically close to the HYFTs initially retrieved in the query sequence; the search space for HYFT extension is limited following a maximum search radius of BLOSUM62 scored substitutions. The results of this extended HYFT-search are shown in Fig. 7, 8 and 9. The extended HYFT-search shows a significant improvement of the average precision, increasing from 0.094 to 0.187. The search is now able to recall 18.9% of all true positives. However, there is also a significant increase in False Positives, as the FPR is going from 3 × 10^*−*5^ to 7 × 10^*−*5^. Because the plots indicate a sharp cutoff point, these False Positives can be filtered out with an appropriate threshold. The distribution of retrieved True Positives is plotted in Fig. 9.

**Table 1.** Average Precision (AP), Mean Average Precision (MAP) and ROC_50_ (area under the ROC curve plotted until 50 TN are found) for both BLAST and HYFT-search.

|  | AP | MAP | ROC<sub>50</sub> |
| --- | --- | --- | --- |
| BLASTP | 0.294 | 0.376 | 0.204 |
| HYFT-search | 0.109 | 0.168 | 0.102 |
| HYFT-search (extended) | 0.187 | 0.261 | 0.160 |

**Fig 7.**
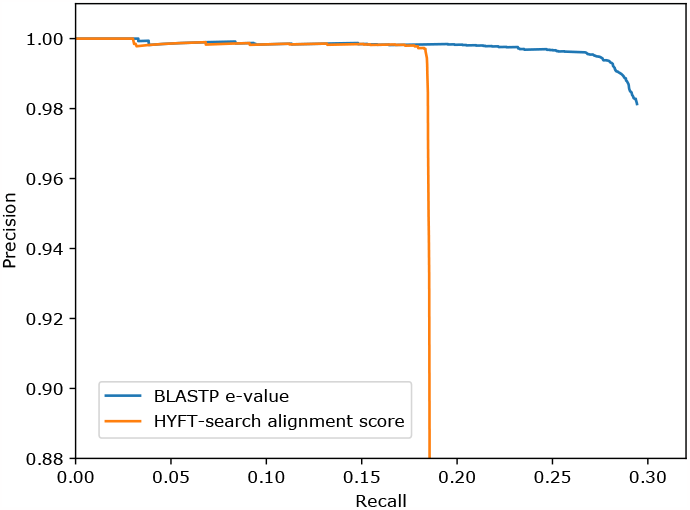
Precision-recall curve for BLASTP and extended HYFT-search, using the e-value (BLASTP) and the alignment score (HYFT-search) for sorting the results.

**Fig 8.**
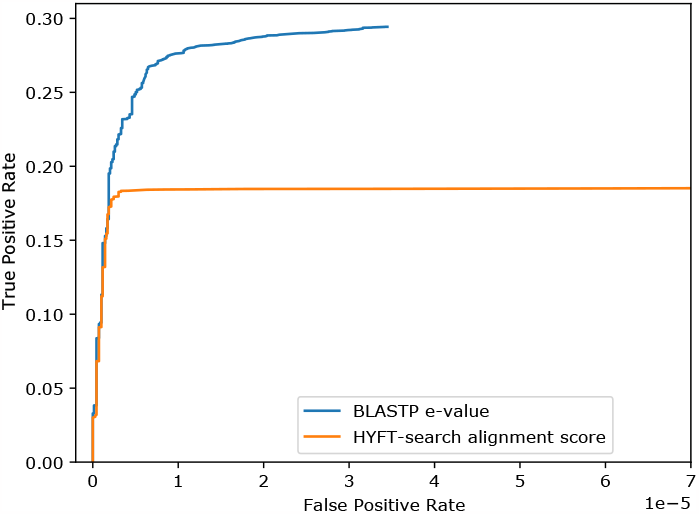
ROC curve for BLASTP and extended HYFT-search, using the e-value (BLASTP) and the alignment score (HYFT-search) for sorting the results.

**Fig 9.**
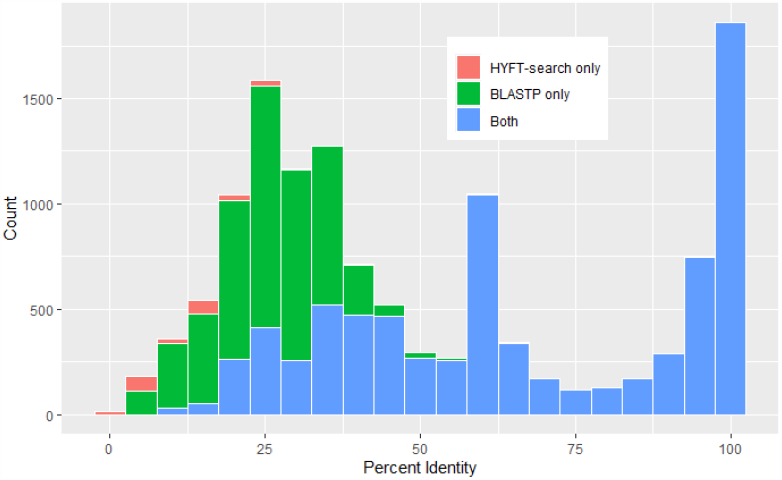
Distribution of True Positives with respect to identity score for BLASTP and extended HYFT-search.

## Discussion

The main purpose of sequence search engine such as BLAST is to retrieve homologous sequences, i.e. sequences sharing a common ancestry through the evolutionary history of life. Homology is typically inferred through sequence alignment, which gives an idea of local sequence similarity. However, it is well known that generally speaking, protein structure tends to be more conserved than sequence [43–45]; hence, sequence similarity metrics are not sufficient to assess whether a query and a result are homologous, specifically toward the lower end of percent identity (*<* 40%) where the signal is blurred [46]. As shown in Fig. 10 approximately 88% of sequences that are structural homologs (expected true positives) have an identity score lower than 50%.

**Fig 10.**
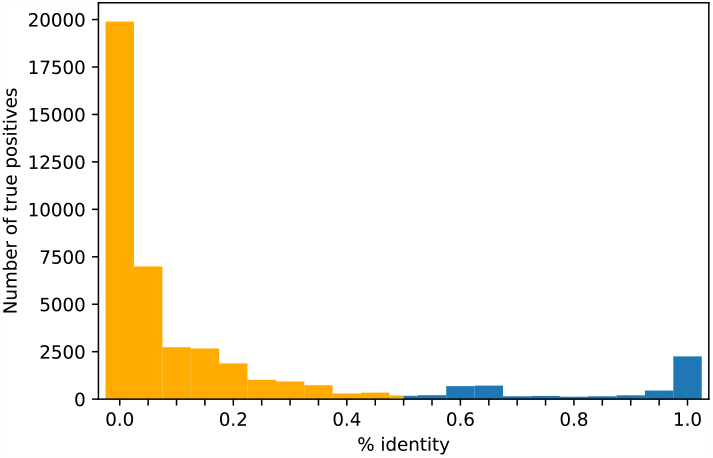
Overall distribution of expected true positives as function of percent identity with respect to query sequences. The results with percent identity below 0.5 are highlighted in orange.

Comparing Figs. 9 and 10 and the scale of the two distributions, it is apparent that the majority of expected true positives (structural homologs) are not found by either methods, highlighting the limit of alignment-based methods for homology inference as was already observed [46] in the past. For the higher ends of the percentage identity scale, sequence similarity should be sufficient to assess homology.

The HYFT-search retrieval of TP decreases below 60 − 65% percent identity. This is explained by the nature of the HYFT parsing approach for indexing. The HYFT parsing relies on non-overlapping HYFTs patterns with the lowest ranking, thus the highest *specificity* with regard to sequence context: specific HYFTs found in a sequence are less likely to be shared by other sequences with diverging identity.

We demonstrated that non exact matching using a word substitution algorithm that creates new HYFTs, which are syntactically close the the HYFTs found in the query, can improve the recall of sequences with percent identity below 65%, albeit at the cost of high FP hits. However, in many use cases such as mapping, automatic annotation, *etc*., the sequences of interest are those with a high similarity to the query. In that case, HYFT-search retrieves the same sequences of interest as the BLASTP method.

The relationship between the pairwise local alignment score (using the Smith-Waterman algorithm [47] with a gap penalty of − 11 and gap extension penalty of − 1) and the percent identity versus the number of common HYFT pattern is illustrated in Figs. 11 and 12. There is an overall trend between the alignment score (respectively identity score) and the number of common non-overlapping, specific HYFTs patterns in pairs, which is expected due to the nature of the metrics. Given these results, we also foresee potential use of HYFTs in other applications benefiting from a parsing-sorting-matching scheme, such as sequence read archiving, multiple sequence alignment, variant calling, downstream analyses and so on.

**Fig 11.**
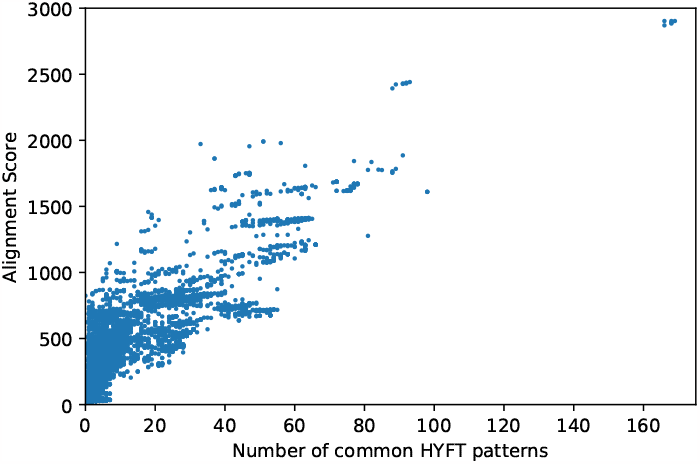
For the BLAST results: local alignment score (gap penalty = − 11, gap extension penalty − = 1) *versus* number of common HYFT pattern with respect to query sequence.

**Fig 12.**
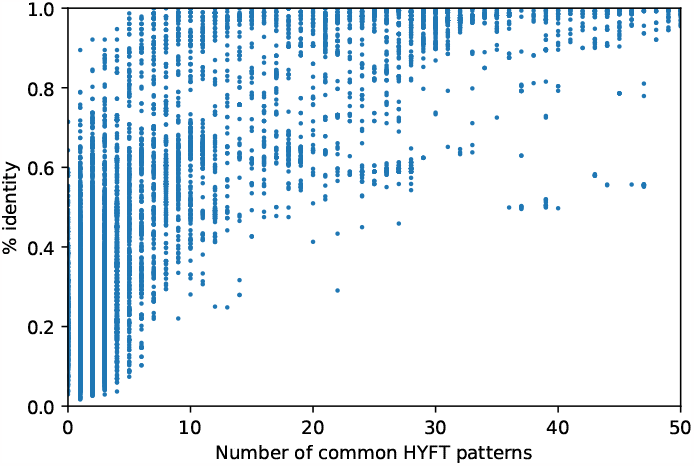
For the BLAST results: percent identity *versus* number of common HYFT pattern with respect to query sequence.

Furthermore, with the ever-increasing availability of biological sequence data, there is a need for scalable methods to retrieve relevant sequences from databases based on similarity. Instead of solving this problem via sequence alignment, the HYFT-based methodology is inspired by NLP methods used for database indexing. Doing so a local sequence alignment is avoided and replaced by a static database lookup of informative patterns. This static indexing results in a system for the retrieval of related sequences that is scalable to databases covering the entire biome. Moreover, removing the need for dynamic programming to compute similarity scores results in quicker turn-around times and the possibility of sparser use of computational resources.

## Conclusions

This benchmarking study comparing the quality of sequence retrieval between BLAST and the HYFT methodology shows that BLAST is able to retrieve more distant homologous sequences with low percent identity than the HYFT-based search. However, when interested in sequences with percent identities over 60%, HYFT-search retrieves the same sequences as BLAST. This makes the HYFT methodology very relevant for use cases where we are interested in highly conserved homologs, such as finding matching paratope regions in antibodies, for example. Additionally, experiments with extending the search using HYFT synonyms to increase the recall show promising results. Furthermore, the HYFT methodology is extremely scalable as it does not rely on sequence alignment to find similars, but uses a parsing-sorting-matching scheme. On this ground, we argue that the HYFT-based indexing is a solution to the biological sequence retrieval in a Big Data context.

## Acknowledgments

We thank V. Akunda for fruitful discussions.

HYFT-based search is currently implemented in the LENSai platform, proprietary software and technology of BioStrand BV. BioStrand BV is an independently operating subsidiary of ImmunoPrecise Antibodies Ltd. LENSai and HYFT are trademarks of BioStrand BV.

The benchmarking study was performed by A.-J. Rousseau, whereas statistics about HYFTs were provided by BioStrand. All authors contributed to the manuscript.

This research received funding from the Flemish Government (Agentschap Innoveren en Ondernemen), HermesFund project number HBC.2020.2182. A.-J. Rousseau received funding from the Flemish Government under the “Onderzoeksprogramma Artificiële Intelligentie (AI) Vlaanderen” program.

